# Diverging Arabidopsis populations quickly accumulate pollen-acting genetic incompatibilities

**DOI:** 10.1101/2024.05.14.594187

**Authors:** Chris Condon, Fantin Carpentier, Marie Tabourin, Natalia Wozniak, Margarita Takou, Christelle Blassiau, Vinod Kumar, Björn Pietzenuk, Rémi Habert, Juliette De Meaux, Ute Krämer, Camille Roux, Russell Corbett-Detig, Vincent Castric

## Abstract

The process by which species diverge from one another, gradually accumulate genetic incompatibilities and eventually reach full-fledged reproductive isolation is a key question in evolutionary biology. However, the nature of reproductive barriers, the pace at which they accumulate and their genomic distribution remain poorly documented. The disruption of co-adapted epistatic interactions in hybrids and the accumulation of selfish genetic elements are proposed contributors to this process, and can lead to the distortion of the mendelian segregation of the affected loci across the genome. In this study we detect and quantify segregation distortion across the genomes of crosses produced from a diverse sampling of *Arabidopsis lyrata* and *A. halleri* populations, two species at the early stages of speciation and that can still interbreed. We show that both the frequency of occurrence and the magnitude of distortion loci increase as the parents’ genetic distance from one another increases. We also observe that distorter loci evolve rapidly, as they occur not only within interspecific hybrids, but also in intraspecific hybrids produced from isolated population crosses. Finally, we identify both genome-wide non-independence and two specific genomic regions on different chromosomes where opposite distortion effects are repeatedly observed across multiple F1 individuals, suggesting negative epistasis is a major contributor to the evolution of hybrid segregation distortion. Our study demonstrates that pollen-acting segregation distortion is ubiquitous, and contributes not only to the ongoing reproductive isolation between *A. halleri* and *A. lyrata*, but also between very recently diverged populations of the same species.

## Introduction

An essential goal of evolutionary biology is to identify the genetic basis and underlying biological processes for speciation. Dobzhansky-Muller incompatibilities (DMIs) are thought to be a major contributor to speciation, as it is a primary explanation for the accumulation of reproductive incompatibilities between isolated or diverging populations (Coyne, 1992; Coyne & Orr, 2004; Dobzhansky, 1982; Muller, 1942). The DMI model hypothesizes that interactions between different loci explains hybrid breakdown. Hybrid genomes contain new combinations of alleles that have not co-evolved together. As a result, most of these new interactions consist of negative epistasis (Simon et al., 2018). The total amount of incompatibilities is expected to depend on the level of molecular divergence between parental genomes, driving the transition from "populations" to "isolated species" (Roux et al., 2016). DMIs may accumulate sufficiently rapidly to produce population-specific incompatibilities, representing the first steps of the speciation process (Bomblies et al., 2007; R. B. Corbett-Detig et al., 2013; Seymour et al., 2019). However, the pace with which incompatibilities accumulate between populations and between species and how they manifest is only poorly understood (Matute & Coyne, 2010; Ravinet et al., 2017).

An early observation in the field (Haldane, 1922; reviewed in Cowell, 2023) is the disproportionate expression of incompatibilities in hemizygous and heterogametic hybrid individuals (Gadau et al., 1999; Turelli & Orr, 1995), suggesting that DMIs that contribute to hybrid sterility and inviability are generally partially recessive (Muller, 1940; Muller, 1942). Therefore, those expressed in haploid tissues (where their expression is not restricted by dominant alleles as would be the case during the diploid phase in a heterozygous individual) should be particularly influential in reproductive isolation. Specifically, factors acting during the haploid phase of the life cycle can directly affect fertility of F1 individuals by compromising proper gamete development, hence potentially representing efficient barriers to gene flow. The haploid phase of the life cycle therefore offers particularly tractable and appealing insights to reveal processes contributing to the formation of species.

Genetic incompatibilities may contribute to gametic inviability in two ways (Illustrated in Fig S1). Under the classical DMI model, as mutations accumulate in diverging populations, each has a chance of generating a negative interaction with alleles that evolve in the other lineage in a hybrid genetic background (Orr, 1996). These interactions can also be interpreted in the context of the evolution of selfish genetic elements that either bias the meiotic process, destroy or disable those gametes that do not inherit them (Burt & Trivers, 2008). Such selfish elements are often coupled with suppressor alleles, masking their effects within populations. When driver and suppressor alleles are uncoupled in hybrids, reactivated drivers can reduce hybrid fertility (*e.g.,* Phadnis and Orr 2009). Either model can result in segregation distortion of incompatible alleles brought about by negative epistasis and distinguishing among them is critical for understanding the contribution of haploid-acting factors to speciation.

Direct gamete sequencing is a promising approach to detecting segregation distortion in the gamete phase, and it does so by bulk sequencing the gametes of an individual and detecting a skew in the ratios of maternal to paternal alleles (Carioscia et al., 2022; Corbett-Detig et al., 2015, 2019). The classical approach to reveal segregation distortion is to focus on interspecific crosses by generating F1 progeny, intercrossing F1s, then generating a large population of F2 individuals (e.g., Fishman & Willis, 2005; Leppälä et al., 2013). This approach is limiting because it is challenging to raise a sufficiently large F2 population to detect loci that distort within F1 individuals (unless their effect is very strong), and because it is usually impossible to distinguish whether segregation distortion is occurring in the gametic phase or in the zygotic phase. Gamete sequencing overcomes these limitations, as it only requires generation of a single F1 hybrid, which eliminates the burden of generating an F2 population, and allows for investigation of a greater variety of F1 cross types. Another advantage is that each gamete (or gametophyte in the case of pollen) is effectively an individual in a population of millions or more, and thus greatly increases the power to detect extremely small distortion effects that would otherwise be impossible to measure. Finally, because gametes are sequenced prior to the zygotic phase, this ensures that any potential segregation distortion is occurring either during or right after meiosis, and involves processes that destroy rather than simply inactivate gametes. Bulk pollen sequencing has been previously used to identify segregation distortion in F1 hybrids derived from crosses between *Arabidopsis lyrata* and *A. halleri* (Corbett-Detig et al., 2019), but leaves important questions unaddressed. *A. lyrata and A. halleri* are two recently diverged species in the Camelineae tribe of the Brassicaceae family that are currently undergoing reproductive isolation from one another (Roux et al., 2011; Wang et al., 2010). Despite this, they are still capable of forming viable and fertile hybrids, and may still be occasionally exchanging genes between their natural populations (Castric et al., 2010; Roux et al. 2011; Novikova et al., 2016). In our previous study (Corbett-Detig et al., 2019), we used bulk gamete sequencing on two F1 hybrids from *A. lyrata* x *A. halleri* crosses and identified segregation distortion on three chromosomes in both hybrids. However, this study did not answer questions related to the accumulation dynamics of segregation distortion, and whether it is a phenomenon that strictly occurs once species have diverged substantially.

Here, we investigate the pace of evolution and effect sizes of segregation distorters by identifying them at two different steps along the speciation process: between populations and between species of the *Arabidopsis* genus. We use gamete sequencing to measure segregation distortion in fourteen unique F1 hybrids, derived from a diverse combination of crosses within and between species in *A. lyrata* and *A. halleri,* encompassing a range of genetic distances between the parental individuals. By using gamete sequencing data from all F1 progeny, and somatic sequencing data from both F1 hybrids and their parents, we show widespread occurrences of segregation distortion loci across a majority of F1 hybrids. We estimate the genomic locations of these loci, and identify antiparallel distortion patterns that are consistent with the DMI rather than the selfish elements model. Distorter frequency and effect sizes increase in relation to the genetic divergence between parents. Our results imply that segregation distortion in the haploid phase acts early during speciation, and is a likely contributor to hybrid infertility.

## Methods

### Plant Sampling and Crosses

Parental plants were collected in natural populations and grown and propagated vegetatively under greenhouse environments either at the university of Cologne (crosses X08, X09, X10, X11 and X12) or at the university of Bochum (crosses X01, X02, X04, X05, X06, X07, X14, Stein et al., 2017). We crossed parental plants by hand pollination and one F1 individual was selected for each cross. The F1s were also grown under their respective greenhouse conditions and brought to flowering. To evaluate whether the distorters revealed in Corbett-Detig et al., (2019) were fixed within species, we chose parents for inter-specific hybrids from geographic origins different from those in Corbett-Detig et al., (2019). For F1s within *A. halleri*, we crossed parents either from close geographic origins or from distant accessions. For F1s within *A. lyrata*, all parents came from accessions that were geographically relatively distant.

Following Corbett-Detig et al., (2019), we collected 20-50 open flowers in 95% EtOH in a 50mL Falcon tube for each F1. The tubes were gently vortexed to suspend pollen, and first centrifuged at low speed to collect flowers in the bottom. We then pipetted the supernatant with pollen in a clean tube, which we centrifuged for 10 minutes at 13,000 rpm. We discarded the supernatant and let pollen pellets dry overnight at room temperature.

### Library Preparation and Sequencing

For *A. halleri* parents, between 50 and 70 mg young leaf tissues were placed in a 1.5-mL polypropylene tube and shock-frozen in liquid nitrogen. A stainless steel metal bead (5 mm diameter) was added per tube, and leaf tissues were homogenized in adapters pre-cooled to -20°C using a Retsch mixer mill (Type MM 300, Retsch, Haan, Germany) for 1 min at 30 Hz. DNA was extracted from this homogenate using the NucleoMag® Plant kit (Macherey Nagel, Düren, Germany) on an epMotion 5075 robot according to the manufacturer’s instructions (Eppendorf, Hamburg, Germany). The quality, quantity, and integrity of DNA samples were assessed on 0.8% (w/v) TAE-agarose gels, with a Nanodrop 2000 spectrophotometer (ThermoFisher Scientific, Darmstadt, Germany) and a Qubit 2.0 Fluorometer (Invitrogen, Darmstadt, Germany) using the dsDNA BR Assay Kit. The Illumina TruSeq DNA PCR-Free Library Prep kit was used for producing libraries on an epMotion 5075 robot according to the manufacturer’s instructions, with assessment of quantity and quality of PCR-free libraries on a Qubit 2.0 Fluorometer and an Agilent 2100 Bioanalyzer using dsDNA HS Assay kit. Libraries passing QC were diluted, pooled and dispatched to Novogene Europe (Cambridge, United Kingdom) for obtaining 150-bp paired-end sequencing reads at a targeted minimum depth of 20x genome coverage using an Illumina NovaSeq 6000 instrument (Illumina, San Diego, USA). Conditions for DNA extraction from leaves of *A. lyrata* parents, library construction and sequencing were described in Takou et al. (2021). For F1 individuals, we extracted genomic DNA from ground leaf tissue using the Macherey Nagel Nucleospin Plant kit. For pollen samples, ten ceramic spheres were added to a tube containing a pollen pellet and crushed with Mp Biomedicals Lysing Matrix D and spun using the Fast prep 24-5G quick prep rotor with the following manual program: 2x(20 seconds at 4.0m/s). Following Corbett-Detig et al., 2019, we isolated genomic DNA from pollen using the Macherey Nagel Nucleospin food kit with the slight modification that we used columns from the Tissu XS kit to accommodate the limited DNA concentrations. We then enzymatically sheared DNA and constructed indexed libraries (both pollen and leaf samples) using the Tecan – NGS Celero NuGen kit using 2 ng of starting material and 8 cycles of PCR. Illumina sequencing of 2x150bp paired-ends fragments was done at GENOSCREEN (Lille, France).

### Data Availability

Sequencing data are located on SRA under accession number PRJNA750331. Sequencing data for crosses X15 and X16, which are interspecific crosses analyzed previously, were from Corbett-Detig et al., 2019, and reanalysed here to ensure consistency.

### Short Read Alignment and Haplotype Phasing

To detect segregation distortion in the gametes of our F1 progeny, we first identified the group of single nucleotide polymorphisms (SNPs) that come from each ancestral haplotype. To do this, we first aligned all F1 and parental short read data to the MN47 *A. lyrata* reference genome (Hu et al., 2011) and jointly-genotyped each individual using the SNPArcher workflow (Mirchandani et al., 2023) using the Senteion-optimized versions of BWA and GATK (Freed et al., 2017). We retained all aligned reads with a MAPQ of 30 or greater, and removed all reads below the 5% and above the 95% depth of coverage quantiles to mitigate against the effects of large structural variants and repeats which might affect allele frequency estimation.

To mitigate the effects of differential and biased mapping of somatic and germline libraries which might result in strongly skewed ancestry ratios, we mapped only the first read in each pair and subsampled the read lengths in each leaf-pollen pair such that both samples have identical read length distributions (see Corbett-Detig et al. 2019). We additionally removed any variant sites within 100 bp of each other. Next, we haplotype-phased each F1 individual using the trio phasing function of WhatsHap (Martin et al., 2016). To do this, we used the variant sites obtained from each parent, constructed a mother-father-child pedigree, and performed trio phasing to separate variant sites into maternal and paternal haplotype blocks. Once phased, we filtered any non-identically phased or genotyped variant sites in each leaf-pollen pair. We additionally performed the entirety of the above mentioned workflow using the *A. halleri* Auby-1 reference genome v1.0 (Pavan et al. in prep), and confirmed that our results are consistent between genomic references (Figure S2).

### Ancestry Ratio Calculations and Estimation of SD Effect Sizes and Loci Positions

To calculate the ancestry ratios for each sample, we used the intersection of retained ancestry informative sites shared between pollen and leaf samples along each chromosome. At each site, we calculated the ancestry ratio as the ratio of maternally-derived read counts to paternally-derived read counts. For both leaf and pollen samples, we plotted the average ancestry ratios in 1,000 SNP windows. We calculated and plotted the ancestry ratio difference as the difference between somatic and germline ancestry ratio for each window.

Using a maximum likelihood framework that models sequencing error, recombination rate and ancestry informative sites, we estimated both the position and effect size of each candidate distorter (see Corbett-Detig *et al*. 2019). Additionally, every variant site along the chromosome is informative to the effect size and location of a distorting locus due to linkage. At each ancestry informative site along the chromosome, we calculated the recombinational distance between our site, p, and a candidate distorting locus, i, in Morgans using the *A. lyrata* recombination map from (Hämälä et al., 2017). We then converted the distance to a probability that a recombination event has occurred between the two positions. Considering also the probability of sequencing error, there are four unique probabilities for mapping an allele A, at site p, given i. Conversely, there are four unique probabilities for mapping the alternate allele a, under the same conditions (See Corbett-Detig et al. 2019 for details). Thus, we estimated the likelihood of a given distortion coefficient, k, at distorting locus i, by calculating the likelihood of the mapped allele frequencies across the entire chromosome.

To estimate the effect size at a given site p, we first estimated the likelihood of k at site p in both the somatic and germline data. Next, using the somatic estimate of k as a null model, we estimated the effect size at site p by calculating the likelihood ratio of germline k to somatic k. We then expanded this procedure to all ancestry informative sites along the chromosome to find the maximum likelihood distorting position. All scripts and programs used for phasing and SD estimation are available from https://github.com/ccondon894/arabidopsis_SD_workflow.

### Confidence Interval Estimation

We estimated the uncertainty around a presumed distorter’s mapping position by bootstrapping with replacement variant sites along a chromosome and rerunning our maximum likelihood framework. We used the 2.5th and 97.5th percentiles as our confidence intervals after 100 bootstrap replicates. To further improve our confidence interval estimates, we selected a subset of samples for chromosomes 4 and 5 whose individual estimated distorter positions are very close to each other. Under the assumption that they are caused by the same underlying genetic element, we treated each sample as an independent event for the same distorter, and computed the joint likelihoods by taking the sum of log-likelihood among crosses for all bootstrap estimates across the selected samples. This approach significantly decreased the widths of estimated confidence intervals while keeping the joint confidence interval within the range of the individual sample intervals.

### Analysis of Segregation Distortion Patterns in Relation to Parental Origin

We performed Fisher’s Exact Tests on multiple cross groups classified by parental species or geographical location, including (i) within-population, (ii) between-population, within-species and (iii) between-species groups. We compared groups to one another and identified groups where there was strong evidence for significantly higher segregation distortion frequency. Using the same groups as the Fisher’s Exact Test, we also tested for significant difference between effect sizes by Mann Whitney-U test.

### Genome-Wide Non-Independence Tests

To test for non-independence between distorting loci on separate chromosomes, we hypothesized that the mean directional effect size (*k*) for each individual across all chromosomes, and the mean across all individuals would be smaller than expected by chance. That is, we expect maternal and paternal distorting alleles to distort in an antiparallel fashion if these effects are consistent with a DMI model. Here, we define distortion in favor of paternal alleles as negative and distortion towards maternal alleles as positive deviations from 0.5. We then constructed a permutation test as follows: (1) Compute the grand mean segregation distortion effect size for all progeny, (2) Randomize the segregation distortion effects across the progeny by conditioning on the number of distorted chromosomes in each individual, (3) Recompute the mean distortion effect for the permuted progeny distribution and record the proportion with a mean effect smaller than or equal to the estimated mean from the true data, (4) Permute 100,000 times.

### Pairwise Chromosomal Interaction Tests

We considered whether there were antiparallel interactions between specific pairs of chromosomes consistent across progeny, such that one chromosome distorts towards the maternal genome and the other chromosome distorts toward the paternal genome, or vice versa. We tested this hypothesis with a permutation test as follows: (1) Count the number of antiparallel distortion effects (i.e., one paternal and one maternal effect) for each chromosome pair across all progeny, (2) Randomly permute the estimated distortion effects across all chromosomes, then recount all chromosome-pair interactions, (3) Repeat for 1,000,000 permutations and record the number of pairs that occur more frequently than the null distribution. To account for the genealogical non-independence caused by some of the crosses sharing particular parents, in these statistical tests we removed crosses with one or more shared parents.

### Gene annotations in the chromosomal intervals

Gene contents of the confidence intervals on chromosome 4 and 5 were retrieved from the annotation of the *A. lyrata* genome assembly (Hu et al., 2011). A list of pollen-related Plant Ontologies (PO) was manually compiled through searches at (https://ontobee.org/ontology/PO) (Dataset S1). PO annotations for the *A. thaliana* genome (po_anatomy_gene_arabidopsis_tair.assoc.gz, po_temporal_gene_arabidopsis_tair.assoc.gz, 2019-07-11) were downloaded from TAIR (https://www.arabidopsis.org/index.jsp). All Arabidopsis gene identifiers associated with pollen-related POs were extracted from both PO annotation lists, and subsequently combined in one list for annotating the gene contents of the chromosome 4 and chromosome 5 confidence intervals (Datasets S2, S3). A complete redundant gene list containing all PO numbers is also provided, with literature references (Dataset S4). Protein-protein interactions were downloaded from TAIR (TairProteinInteraction.20090527.txt, 2019-07-11).

## Results and Discussion

### Sequencing and Phasing

We obtained an average of 29 million reads for the parental individuals, corresponding to an expected sequencing depth of 41X and providing substantial power to identify SNPs distinguishing parental genomes. For the F1 samples, we obtained an average of 100 million pollen reads and 98 million leaf reads, corresponding to an expected sequencing depth of 144X, allowing for a precise estimation of the parental contribution at each SNP. After mapping all pollen and leaf sequence data to the *A. lyrata* genome, trio phasing, and filtering inconsistently genotyped or phased sites, we obtained an average of 848,054 retained sites shared between pollen and leaf, with hybrid X06 retaining a low of 458,629 sites, and hybrid X15 retaining a high of 1,487,203 sites (Table S2).

### Signatures of Segregation Distortion via Raw Ancestry Ratios

We computed the ancestry ratios across all chromosomes for each hybrid for both somatic and germline read data (Fig. 1, Fig S1). The ancestry ratio is defined as the ratio of reads from the maternal haplotype versus reads from the paternal haplotype for a given site. The somatic genome sequence data provides a null model, and accordingly across each chromosome, the raw somatic ancestry ratios tend to stay near to 0.5, consistent with our expectations of 50:50 representation of haplotypes in somatic tissues. However, in the majority of F1s, and across numerous chromosomes, we observe signatures of segregation distortion in the pollen samples, with skews in both the maternal and paternal directions (Table 1, Fig. 1, Fig. S1). From the somatic and germline ancestry ratios, we calculate the ancestry ratio difference between somatic and pollen samples (Fig. 2). We observe a skew in germline ancestry ratios at least once on every chromosome, and ancestry ratio differences vary dramatically (0–19%).

**Figure 1:**
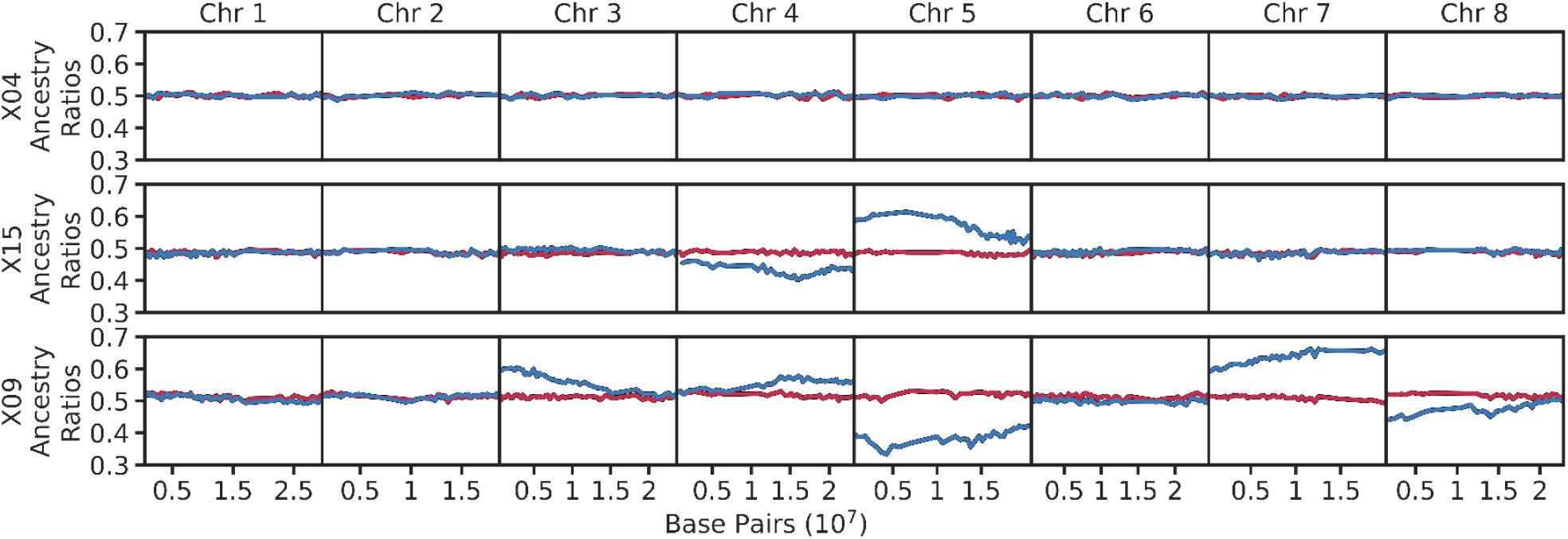
Raw ancestry ratios plotted across all chromosomes for three selected individuals. Included are X04 (intraspecific *A. halleri*), X15 (interspecific) and X09 (interspecific). Ancestry ratios are plotted from somatic (red) and germline (blue) read data in 1000 SNP non-overlapping windows.

**Figure 2:**
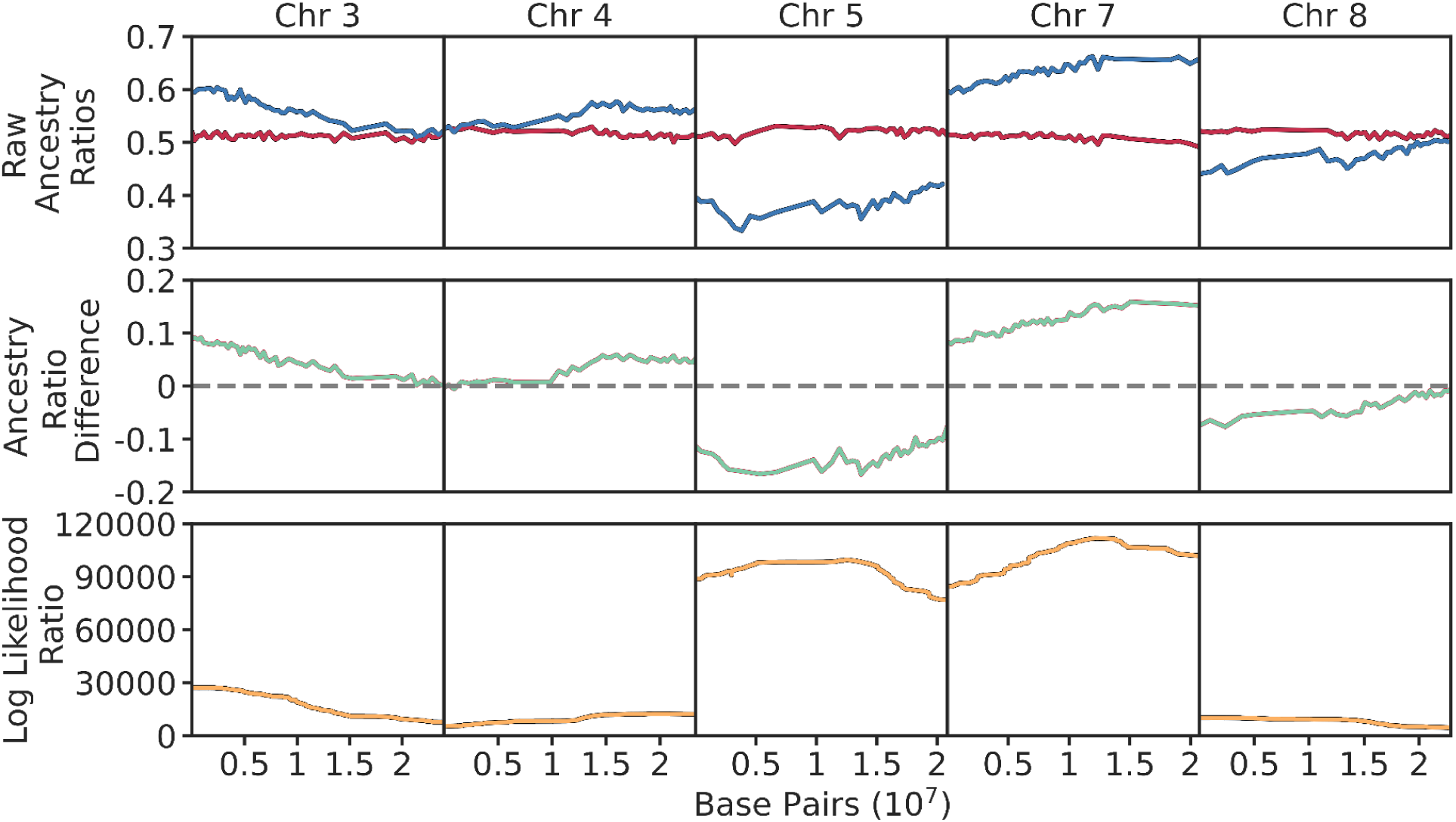
Raw ancestry ratios, ancestry ratio differences, and estimated distorter positions for select chromosomes for interspecific hybrid X09. Raw ratios are plotted from somatic (red) and germline (blue) read data in 1000 SNP non-overlapping windows. Differences between somatic and germline ancestry ratios are colored green. The dashed gray horizontal line indicates the expected difference, zero, when both ratios are equal. The estimated positions of distorting loci along each chromosome are colored orange.

**Table 1:** Table of all F1 hybrids included in this study. Included within this table are distortion effect sizes within each chromosome. Effect sizes are colored dependent on their effect direction towards either the maternal (red) or paternal (blue) genome. Additional information about the parents and populations contributing to each cross are available in Table S2.

| Cross Type | Cross ID | Estimated Distortion Effect Size (k) |  |  |  |  |  |  |  |
| --- | --- | --- | --- | --- | --- | --- | --- | --- | --- |
|  |  | Chr1 | Chr2 | Chr3 | Chr4 | Chr5 | Chr6 | Chr7 | Chr8 |
| Within Population | X04 | 0 | 0 | 0 | 0 | 0 | 0 | 0 | 0 |
|  | X05 | 0 | 0 | 0 | 0 | 0 | 0 | 0 | 0 |
|  | X06 | 0 | 0 | 0 | 0 | 0 | 0 | 0 | 0 |
|  | X07 | 0 | 0 | 0 | 0 | 0 | 0 | 0 | 0 |
| Between Population | X01 | 0 | 0 | 0 | -0.07 | 0.06 | 0 | 0 | 0 |
|  | X02 | 0 | 0 | 0 | 0.07 | -0.05 | 0 | 0 | 0 |
|  | X12 | -0.05 | 0 | 0.1 | 0 | 0 | 0.04 | -0.02 | 0 |
|  | X10 | -0.02 | 0 | 0.02 | 0 | -0.02 | 0 | 0 | 0 |
|  | X11 | 0 | 0 | 0 | 0 | 0 | 0 | 0 | 0 |
| Between Species | X09 | 0 | 0 | 0.1 | 0.06 | -0.19 | -0.02 | 0.19 | -0.08 |
|  | X08 | 0 | 0 | -0.02 | 0 | 0 | 0.02 | 0 | 0 |
|  | X15 | 0 | 0 | 0.02 | -0.08 | 0.13 | 0 | 0 | 0 |
|  | X16 | 0.02 | 0.02 | 0.03 | -0.05 | 0.06 | 0 | 0 | 0 |
|  | X14 | -0.02 | 0 | 0 | -0.02 | 0 | 0 | 0 | 0 |

### Segregation Distortion Patterns Vary Greatly Amongst Different Hybrid Groups

We detected many instances of pollen-acting segregation distortion in crosses both within and between species. We previously developed a maximum likelihood framework to estimate the effect size and position of a putative distorter locus along a chromosome (Corbett-Detig et al., 2019). Using this framework, we estimated both segregation distorter effect size and position along each chromosome for all hybrids, and identified twenty five distorter loci with effect sizes greater than or equal to k=0.02 (Fig. 2). As expected, many distorter loci appear in interspecific hybrids derived from crosses between *A. lyrata* and *A. halleri* parents, though the presence of distorter loci are not restricted to interspecific hybrids, and are also observed in several intraspecific *A. lyrata* or *A. halleri* hybrids.

Chromosome-specific distortion patterns vary and depend on the type of cross (Fig 1). In intraspecific *halleri,* we observe distortion in just two crosses, X01 and X02. This pair shares one parent, Pais09, which is the mother of X01 and the father of X02. Both crosses exhibit similar distorter loci on chromosomes 4 and 5, but each distorter skews in the opposite parental direction. Contrary to other documented cases e.g. in flour beetles (Beeman et al., 1992), mouse (Weichenhan et al., 1996), *Caenorhabditis elegans* (Ben-David et al., 2017) or *C. tropicalis* (Noble et al., 2021), the distortion patterns in this pair suggest that these distorters do not exhibit a maternal effect, because the ancestry ratio is associated with male or female contribution. The remaining four *halleri* hybrids do not show distortion (Table 1). Notably, all are derived from crosses whose parents are from nearby geographic locations. Genetic differences between parents in X01 and X02 is slightly higher than between those parents in non-distorting *halleri* hybrids (π = 0.0052 vs. π = 0.0043-0.0049, see Table S1). Although they are the same species, our results suggest that the ancestral populations of X01 and X02 have accumulated sufficient genetic incompatibilities that they are readily measurable within the power of our experimental design.

Intraspecific *A. lyrata* crosses display distortion patterns distinct from intraspecific *halleri* crosses. All three have a mother from Norway and a father from Germany. Two crosses demonstrate distortion patterns on chromosomes 1 and 3 in the same parental directions. However, X12 shows stronger signatures of distortion genome-wide. Chromosomes 1 and 3 have much larger effect sizes, and X12 also exhibits distortion on chromosomes 6 and 7, while X10 contains a unique distorter locus on chromosome 5. These differences cannot be explained by parental genetic divergence, since all three have similar levels of genetic differentiation and are derived from the same set of populations (Table S1). Instead, these results demonstrate a level of variability in the presence or penetrance of distorting loci. This variability could be caused by the presence of modifiers across the genome modulating expression of the distorters, by allelic variability of the elements causing the distortion in the different individuals, or by other sources of individual variation in phenotypic expression, such as environmental effects.

The number of distorter loci and effect sizes in interspecific crosses exceeds those found in intraspecific crosses. Of the 29 total distorter loci we identified, 18 are in interspecific hybrids, and a distorter locus appears at least once on every chromosome within this group. Effect sizes vary substantially within this group (range |k|=0.02–0.19). We replicated previously observed distortion signals in hybrids X15 and X16 (Corbett-Detig et al. 2019). Additionally, here we report two previously undetected distorter loci on chromosomes 1 and 2 in X16. These additional loci could be explained by the increased statistical power we gained from trio phasing (which is new as compared to our previous analysis), which almost doubled the number of ancestry informative sites retained. X14 and X08 show modest signatures of segregation distortion, with only two loci identified in each hybrid and an effect size of |k|=0.02. X09 however, exhibits distorter loci on six of its eight chromosomes, and two loci display effect sizes of |k|=0.19, by far the most distorter loci and largest effect sizes of all the crosses. Notably, X08 and X09 share a parent, Hall2.2, yet show very contrasting distortion patterns. This reaffirms our observations that not only is segregation distortion evolving rapidly, but occurrences and effect sizes vary substantially.

### Estimated Segregation Distortion Frequency and Effect Size Increase with Genetic Divergence Between Parents

Regardless of the proximal mechanism, a prediction is that segregation distortion should be more frequent in crosses with more genetically divergent parents. To formally test this, we grouped our hybrid progeny into three groups: within-population (n=4), between-populations (n=5), and between-species (n=5). Collectively, within-population and between-population also comprise the “within-species” group. Segregation distortion does not occur in any of the within-population crosses. Segregation distortion is apparent in both between-population groups and between-species groups. When we compare occurrences between within-species and between-species groups, we find that there are significantly more occurrences in the between-species group (Fisher’s Exact Test, P=0.0013). While the difference between the between-species and between-population groups is not significant (FET, P=0.16), the odds ratio of 2.15 suggests that segregation distortion occurs more frequently in the between-species group. Overall, the proportion of distorting chromosomes increases with parental genetic divergence.

A second prediction is that effect sizes (*i.e.,* the proportion of affected pollen) should become larger as parental genetic divergence increases. In comparing between-species to within-species groups (Fig. 3), we find the effect sizes are significantly higher (Mann-Whitney U, P=0.0006). There is no significant difference in effect size when comparing between-species to between-population groups (Mann Whitney U, P=0.10), but the mean distortion effect size observed in between-species crosses is larger. Overall, these results demonstrate that distortion patterns are not limited to genomic differences between *A. lyrata* and *A. halleri*, because we observe them in many between-population crosses. Hence, parental genetic distance is a predictor of the distribution and abundance of distortion effects.

**Figure 3:**
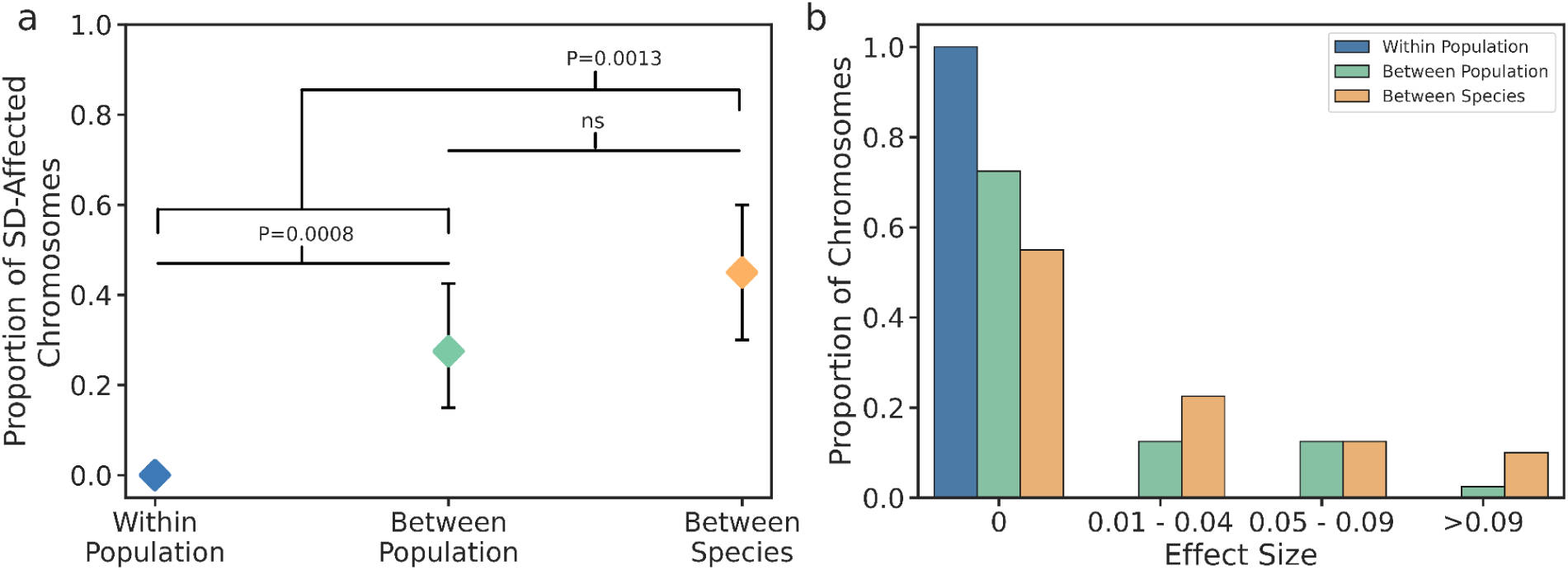
(**a**) Proportion of segregation distortion presence/absences in chromosomes sorted by cross group. (**b**) Proportion of segregation distortion effect sizes, sorted by cross group and effect size cluster.

### Evidence for Non-Independence Among Loci

If negative interactions between parental genomes (DMI) are the source of most segregation distortion, we expect that the effects of maternal and paternal alleles will often be paired by an apparent distortion effect from the opposite parental genome. In contrast, if selfish elements are causing the distortion, we expect independent or directional distortion effects (see Fig S1). To distinguish between these two possibilities, we tested whether the sum of k for a given F1 was closer to 0 than we expect by chance (as expected under the DMI model). In evaluating this hypothesis, we found that there is a significant correlation between maternal-paternal directional effect size such that the sum of all significant effects in a cross is closer to zero than expected by chance (P=0.001, Permutation Test, Fig. 4a). This suggests that interactions among alleles contributed by the maternal and paternal genomes are a major contributor to pollen inviability.

**Figure 4:**
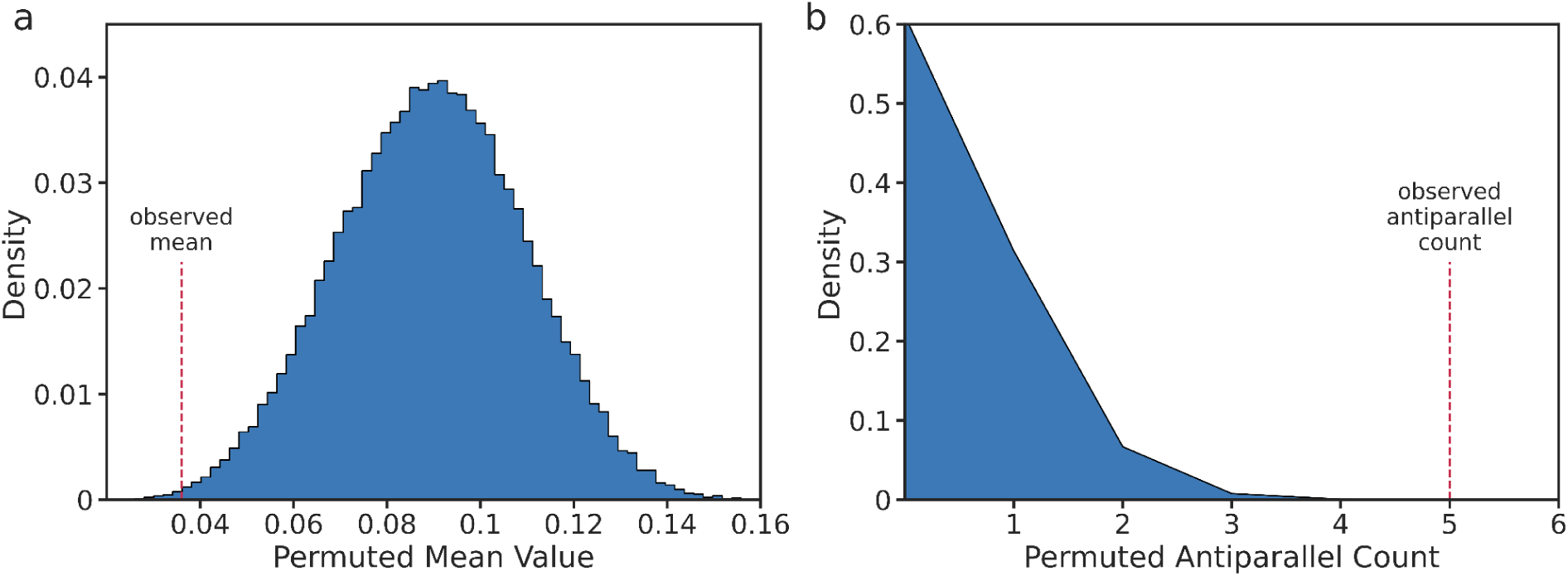
(**a**) The distribution of mean effect sizes for 100,000 permutations testing for non-independence across all F1 progeny. The original mean effect size for our F1 progeny is indicated by the dashed red line. (**b**) The distribution of antiparallel interaction occurrences between chromosomes 4 and 5 for 1,000,000 permutations. The original antiparallel count is indicated by the dashed red line.

Next, we considered whether there were non-independent, antiparallel interactions between specific pairs of chromosomes consistent across progeny. We found that the pair of chromosomes 4 and 5 show highly significant non-independence across the set of crosses (Bonferroni adj., P=0.00005, Permutation Test, Fig. 4b), suggesting a genetic interaction between this specific pair of chromosomes. This further supports the idea that interactions between alleles contributed by each parental genomes is the major contributor to the pollen-acting segregation distortion. We note that to fully take into account phylogenetic non-independence in these tests (such as *e.g.* the fact that two *A. halleri* individuals are more likely to contain the same distorting allele rather than each being an independent evolutionary process generating the observation) would require experimental crosses to control in a more precise manner the genealogy of the distorting chromosomes being observed.

### Confidence Interval Estimation Around Distorting Loci on Chromosomes 4 and 5

Next we estimated the uncertainty around a putative distorter’s mapping position by bootstrapping, as in Corbett-Detig et al. (2019). To improve on confidence interval estimates from individual crosses, we selected a subset of hybrid samples for chromosomes 4 and 5 whose individual estimated distorter positions are close enough that they could presumably correspond to the same locus. Following this logic, we proposed that each sample could be treated as an independent event for the same distorter, and thus computed the joint likelihoods for all bootstraps across the selected samples (Fig. 5). This approach greatly improved the confidence interval estimates compared to the analysis based on single crosses, decreasing the window size while also keeping the joint confidence interval within the range of the individual sample intervals (more so for chromosome 4 than chromosome 5). The final joint confidence interval sizes for chromosomes 4 and 5 were 2.14 Mbp (17,065,274 – 19,205,774 bp) and 610 kbp (4,032,300 – 4,643,400 bp), respectively, on the *A. lyrata* reference genome. We acknowledge that this approach is only valid assuming that distorters in close proximity to each other are indeed the same distorting allele and that no secondary distorters are present. Thus, these joint confidence intervals should be interpreted cautiously, and any further examination of these distorter positions would require functional follow-up work.

**Figure 5:**
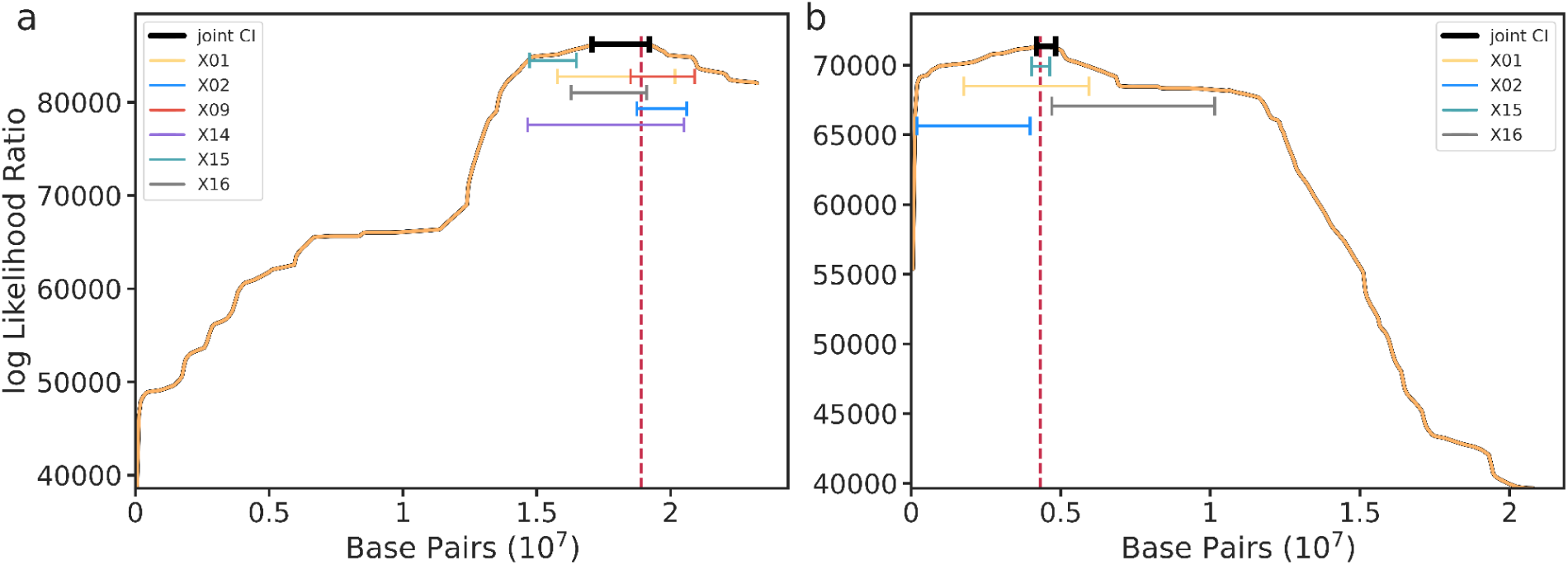
Joint-likelihood estimates and confidence intervals of distorting loci for chromosome 4 (**a**) and chromosome 5 (**b**). The solid orange line indicates the estimated likelihood of the distorting locus along the chromosome. The dashed red line indicates the joint maximum-likelihood position and solid black bars indicate the joint confidence interval. Colored bars indicate the individually estimated sample confidence intervals.

### Candidate Segregation Distortion Genes

The joint chromosome 4 and chromosome 5 confidence intervals were predicted to contain 411 and 61 genes, respectively in the *A. lyrata* reference genome (Datasets S2, S3)(Hu et al., 2011). Among them, 216 and 26 had *A. thaliana* orthologs annotated as pollen-related in the Plant Ontology (PO) database (www.plantontology.org, see list of PO terms in Dataset S1). The predicted functions of a number of these genes could be relevant in the context of the observed segregation distortion phenotype, such as genes encoding F-box proteins, proteases, proteins contributing to transcription and translation, cell division, nutrient and metabolite transport, endomembrane trafficking and signaling. For instance, the chromosome 5 interval contains a gene encoding a Concanavalin A-like lectin-type protein kinase family protein (AL5G17300) in syntenic position with At5G65600 (AtLecRK-IX.2). AtLEcRK-IX.2 was characterized as a regulator of cell death and a positive regulator of pathogen recognition receptor-triggered immunity in Arabidopsis acting to phosphorylate RbohD and thus triggering ROS production (Luo et al., 2017; Wang et al., 2015). A gene within the chromosome 4 confidence interval (AL4G34480) is a non-syntenic homologue of AT3G56440 encoding AtATG18D, an uncharacterized protein related to yeast autophagy 18 (ATG18) D (Xiong et al., 2005). These are examples for which an alteration of the activity of gene products could trigger cell death during pollen development. None of the gene products in the chromosome 4 interval were previously identified to establish direct protein-protein interactions with the products of any of the genes in the chromosome 5 interval according to the protein-protein interactions available through TAIR, and none of the genes in the two intervals had previously been identified as being required for male gametophyte development (Muralla et al., 2011). We acknowledge that the sheer number of possible direct or indirect molecular interactions between any of these genes currently makes it difficult to formally pinpoint specific pairs of genes more precisely and that one or both causal genes might be absent in the confidence intervals of the reference genome used here. Hence, our method serves as an efficient way to narrow down a segregation distortion locus to a limited chromosomal interval, but fine-mapping will now be necessary to further restrict the list of candidate genes.

## Conclusion

Our results demonstrate that segregation distortion is widespread and rapidly evolving not only between different *Arabidopsis* species, but within species as well. A rapid spread of segregation distorters has been observed in several instances (Seymour et al. 2019), but is often only transient and followed by the rapid evolution of suppression (reviewed in Price et al., 2019). Here we find that segregation distortion is highly variable among individuals, but the general trend is that it increases in frequency and magnitude as parental lineages diverge. This approach is straightforward to apply to many other species, and expanding the analysis to more species will now allow to determine the generality of this conclusion. An especially interesting comparison would be to analyze hybrids between selfing vs outcrossing lineages, as differences in the mating system are expected to lead to differences in the intensity of genetic conflicts (Brandvain & Haig, 2005). We expect that the dynamics of driver-suppressors should be more apparent in such circumstances.

More generally, our results demonstrate that segregation distortion acts early during population divergence and may be a contributor towards the common pattern of hybrid infertility contributing to speciation. We cannot completely rule out the driver-suppressor model, but the pattern of segregation distortion observed in our data, where skews in parental directions tend to compensate each other both overall and at specific loci, is more consistent with negative epistatic interactions between pairs of loci, where the viability of the gametes that contain specific combination of alleles is impaired. In particular, it will be exciting to determine how different groups of organisms accumulate such incompatibilities, especially since the “speciation clock” has been suggested to tick at different paces across the tree of life (e.g. in plants vs animals, Monnet et al., 2023). In addition, here we focused on the male germline, but the comparison with the female germline would be necessary to provide a comprehensive picture.

## Supplementary

**Figure S1:**
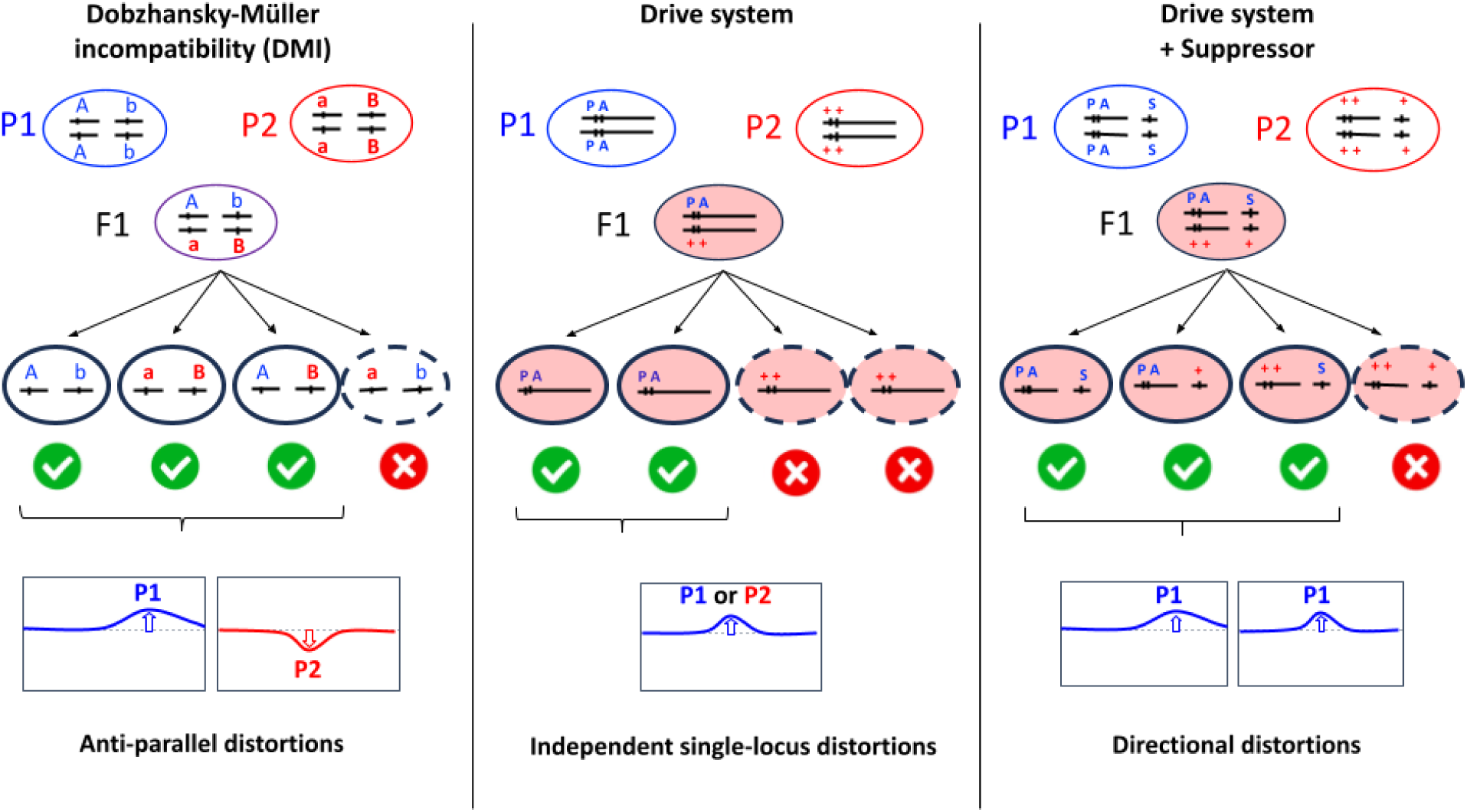
Illustration of three possible models of segregation distortion. **A.** The DMI model illustrates gametophytic incompatibility between alleles a and b (derived alleles, independently fixed in either parental species), resulting in antiparallel distortions towards alleles from P1 and from P2 between the two loci. **B.** In drive systems, a toxic element is expressed at the diploid stage, and only gametic products inheriting the antidote survive. All drive systems documented rely on genetic elements that are tightly linked (either a Poison-Antidote, PA system as represented here, or a Killer-Target system where a “killer” element is produced by one chromosome selectively kills chromosomes carrying a “target” element in *trans*). Such single-locus drive systems are expected to result in independent distortion patterns across chromosomes carrying them (either towards P1 or towards P2, but independently from one another). **C.** Suppressors (S) of drive systems are commonly found in natural populations, and are often genetically unlinked to the original driver elements. In such a case, the suppressor would rescue gametes carrying it, resulting in distortion towards the parent contributing to the driver and the suppressor.

Figure S2: Comparison of ancestry ratio results for each cross hybrid using *A. lyrata* and *A. halleri* reference genomes. Ancestry ratios are plotted from somatic (red) and germline (blue) read data in 1000 SNP non-overlapping windows.

(will be attached as PDF)

**Table S1.**
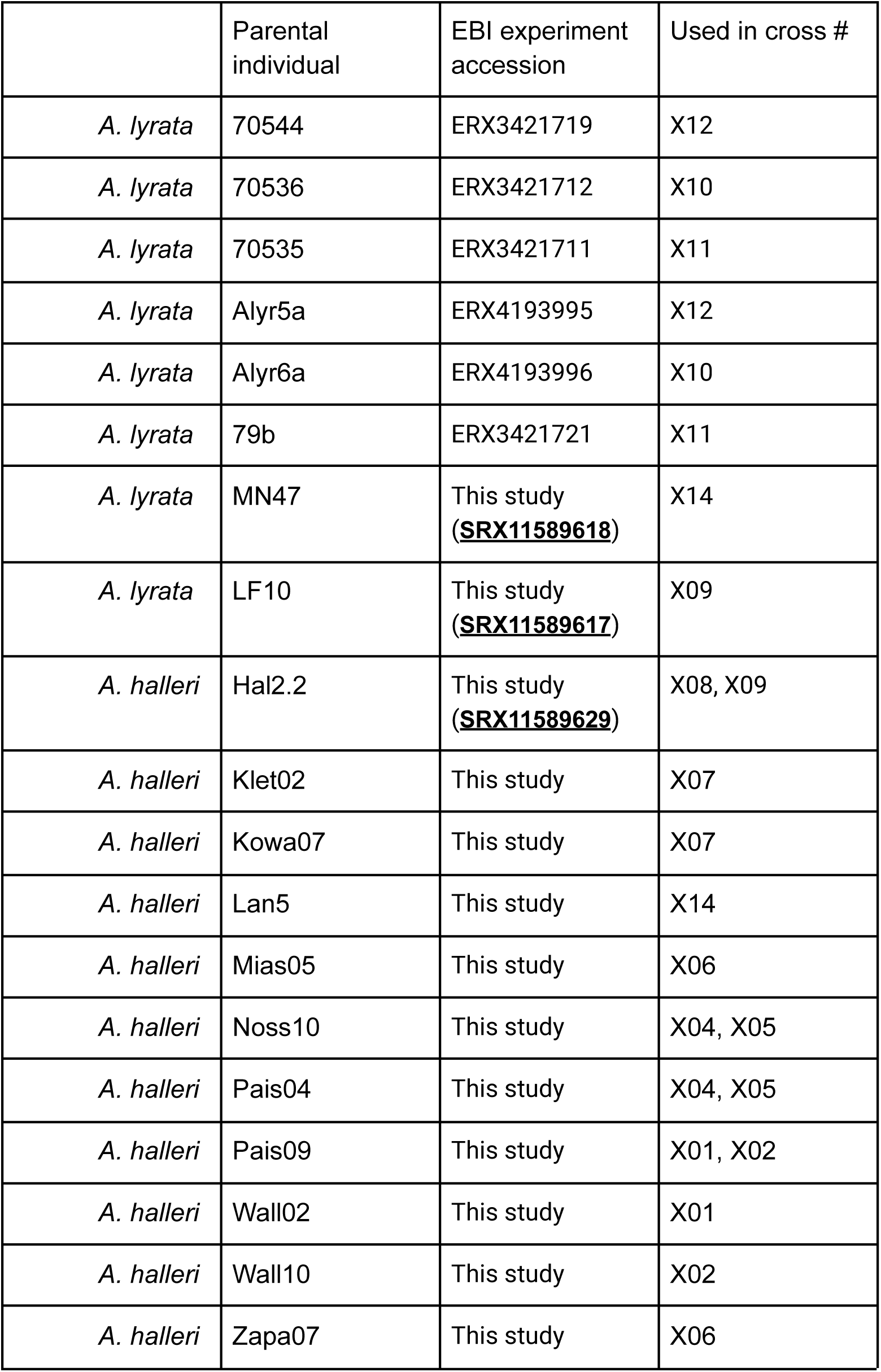
Origin of the sequencing reads for parental individuals used in this study.

|  | Parental individual | EBI experiment accession | Used in cross # |
| --- | --- | --- | --- |
| *A. lyrata* | 70544 | ERX3421719 | X12 |
| *A. lyrata* | 70536 | ERX3421712 | X10 |
| *A. lyrata* | 70535 | ERX3421711 | X11 |
| *A. lyrata* | Alyr5a | ERX4193995 | X12 |
| *A. lyrata* | Alyr6a | ERX4193996 | X10 |
| *A. lyrata* | 79b | ERX3421721 | X11 |
| *A. lyrata* | MN47 | This study<br>(**<u>SRX11589618</u>**) | X14 |
| *A. lyrata* | LF10 | This study<br>(**<u>SRX11589617</u>**) | X09 |
| *A. halleri* | Hal2.2 | This study<br>(**<u>SRX11589629</u>**) | X08, X09 |
| *A. halleri* | Klet02 | This study | X07 |
| *A. halleri* | Kowa07 | This study | X07 |
| *A. halleri* | Lan5 | This study | X14 |
| *A. halleri* | Mias05 | This study | X06 |
| *A. halleri* | Noss10 | This study | X04, X05 |
| *A. halleri* | Pais04 | This study | X04, X05 |
| *A. halleri* | Pais09 | This study | X01, X02 |
| *A. halleri* | Wall02 | This study | X01 |
| *A. halleri* | Wall10 | This study | X02 |
| *A. halleri* | Zapa07 | This study | X06 |

**Table S2:** Table of all F1 hybrids included in this study. Included within this table are distortion effect sizes within each chromosome, species, cross type, mother, father and parental genetic distance calculated as pi. Effect sizes are colored dependent on their effect direction towards either the maternal (red) or paternal (blue) genome.

| Cross | Species | Cross Type | Population Type | Mother | Father | Parental Genetic Distance | Chr1 Effect Size | Chr2 Effect Size | Chr3 Effect Size | Chr4 Effect Size | Chr5 Effect Size | Chr6 Effect Size | Chr7 Effect Size | Chr8 Effect Size |
| --- | --- | --- | --- | --- | --- | --- | --- | --- | --- | --- | --- | --- | --- | --- |
| X06 | halleri | within species | Within | Mias05 (Poland) | Zapa07 (Poland) | 0.0043 | 0 | 0 | 0 | 0 | 0 | 0 | 0 | 0 |
| X07 | halleri | within species | Within | Klet02 (Poland) | Kowa07 (Poland) | 0.0043 | 0 | 0 | 0 | 0 | 0 | 0 | 0 | 0 |
| X04 | halleri | within species | Within | Noss10 (Italy) | Pais04 (Italy) | 0.0049 | 0 | 0 | 0 | 0 | 0 | 0 | 0 | 0 |
| X05 | halleri | within species | Within | Pais04 (Italy) | Noss10 (Italy) | 0.0049 | 0 | 0 | 0 | 0 | 0 | 0 | 0 | 0 |
| X01 | halleri | within species | Between | Pais09 (Italy) | Wall02 (Germany) | 0.0052 | 0 | 0 | 0 | -0.07 | 0.06 | 0 | 0 | 0 |
| X02 | halleri | within species | Between | Wall10 (Germany) | Pais09 (Italy) | 0.0052 | 0 | 0 | 0 | 0.07 | -0.05 | 0 | 0 | 0 |
| X12 | lyrata | within species | Between | 70544 (Norway) | Alyr5a (Germany) | 0.0052 | -0.05 | 0 | 0.1 | 0 | 0 | 0.04 | -0.02 | 0 |
| X10 | lyrata | within species | Between | 70536 (Norway) | Alyr6a (Germany) | 0.0054 | -0.02 | 0 | 0.02 | 0 | -0.02 | 0 | 0 | 0 |
| X11 | lyrata | within species | Between | 70535 (Norway) | 79b (Germany) | 0.0055 | 0 | 0 | 0 | 0 | 0 | 0 | 0 | 0 |
| X09 | hybrid | between species | N/A | Al: LF10 (Austria) | Ah: Hall2.2 (Italy) | 0.008 | 0 | 0 | 0.1 | 0.06 | -0.19 | -0.02 | 0.19 | -0.08 |
| X08 | hybrid | between species | N/A | Al: 79b (Germany) | Ah: Hall2.2 (Italy) | 0.0084 | 0 | 0 | -0.02 | 0 | 0 | 0.02 | 0 | 0 |
| X15 | hybrid | between species | N/A | Al: CP1-1 (Czech rep.) | Ah: l14 (Italy) | 0.0087 | 0 | 0 | 0.02 | -0.08 | 0.13 | 0 | 0 | 0 |
| X16 | hybrid | between species | N/A | Al: CP991 (Czech rep.) | Ah: l16 (Italy) | 0.0088 | 0.02 | 0.02 | 0.03 | -0.05 | 0.06 | 0 | 0 | 0 |
| X14 | hybrid | between species | N/A | Al: MN (USA) | Ah: Lan5 (Germany) | 0.009 | -0.02 | 0 | 0 | -0.02 | 0 | 0 | 0 | 0 |

## References

Beeman, R. W., Friesen, K. S., & Denell, R. E. (1992). Maternal-effect selfish genes in flour beetles. *Science (New York*, N.Y.), 256(5053), 89–92. 10.1126/science.1566060

Ben-David, E., Burga, A., & Kruglyak, L. (2017). A maternal-effect selfish genetic element in Caenorhabditis elegans. *Science (New York*, N.Y.), 356(6342), 1051–1055. 10.1126/science.aan0621

Bomblies, K., Lempe, J., Epple, P., Warthmann, N., Lanz, C., Dangl, J. L., & Weigel, D. (2007). Autoimmune Response as a Mechanism for a Dobzhansky-Muller-Type Incompatibility Syndrome in Plants. PLOS Biology, 5(9), e236. 10.1371/journal.pbio.0050236

Brandvain, Y., & Haig, D. (2005). Divergent mating systems and parental conflict as a barrier to hybridization in flowering plants. The American Naturalist, 166(3), 330–338. 10.1086/432036

Burt, A., & Trivers, R. (2008). Genes in Conflict: The Biology of Selfish Genetic Elements. Belknap Press.

Carioscia, S. A., Weaver, K. J., Bortvin, A. N., Pan, H., Ariad, D., Bell, A. D., & McCoy, R. C. (2022). A method for low-coverage single-gamete sequence analysis demonstrates adherence to Mendel’s first law across a large sample of human sperm. eLife, 11, e76383. 10.7554/eLife.76383

Castric, V., Bechsgaard, J. S., Grenier, S., Noureddine, R., Schierup, M. H., & Vekemans, X. (2010). Molecular evolution within and between self-incompatibility specificities. Molecular Biology and Evolution, 27(1), 11–20. 10.1093/molbev/msp224

Castric, V., Bechsgaard, J., Schierup, M. H., & Vekemans, X. (2008). Repeated adaptive introgression at a gene under multiallelic balancing selection. PLoS Genetics, 4(8), e1000168. 10.1371/journal.pgen.1000168

Corbett-Detig, R. B., Zhou, J., Clark, A. G., Hartl, D. L., & Ayroles, J. F. (2013). Genetic incompatibilities are widespread within species. Nature, 504(7478), 135–137. 10.1038/nature12678

Corbett-Detig, R., Jacobs-Palmer, E., Hartl, D., & Hoekstra, H. (2015). Direct Gamete Sequencing Reveals No Evidence for Segregation Distortion in House Mouse Hybrids. PloS One, 10(6), e0131933. 10.1371/journal.pone.0131933

Corbett-Detig, R., Medina, P., Frérot, H., Blassiau, C., & Castric, V. (2019). Bulk pollen sequencing reveals rapid evolution of segregation distortion in the male germline of Arabidopsis hybrids. Evolution Letters, 3(1), 93–103. 10.1002/evl3.96

Cowell, F. (2023). 100 years of Haldane’s rule. Journal of Evolutionary Biology, 36(2), 337–346. 10.1111/jeb.14112

Coyne, J. A. (1992). Genetics and speciation. Nature, 355(6360), Article 6360. 10.1038/355511a0

Coyne, J. A., & Orr, H. A. (2004). Speciation. Sinauer Associates. http://catdir.loc.gov/catdir/toc/ecip0417/2004009505.html

Dobzhansky, T. (1982). Genetics and the Origin of Species: Columbia Classics edition (p. 364 Pages). Columbia University Press.

Fishman, L., & Willis, J. H. (2005). A novel meiotic drive locus almost completely distorts segregation in mimulus (monkeyflower) hybrids. Genetics, 169(1), 347–353. 10.1534/genetics.104.032789

Freed, D., Aldana, R., Weber, J., & Edwards, J. (2017). The Sentieon Genomics Tools—A fast and accurate solution to variant calling from next-generation sequence dat. 10.1101/115717

Gadau, J., Page, R. E., & Werren, J. H. (1999). Mapping of hybrid incompatibility loci in Nasonia. Genetics, 153(4), 1731–1741. 10.1093/genetics/153.4.1731

Haldane, J. (1922). Sex ratio and aunisexual sterility in hybrid animals. Journal of Genetics, 12, 101–109.

Hämälä, T., Mattila, T. M., Leinonen, P. H., Kuittinen, H., & Savolainen, O. (2017). Role of seed germination in adaptation and reproductive isolation in Arabidopsis lyrata. Molecular Ecology, 26(13), 3484–3496. 10.1111/mec.14135

Hu, T. T., Pattyn, P., Bakker, E. G., Cao, J., Cheng, J.-F., Clark, R. M., Fahlgren, N., Fawcett, J. A., Grimwood, J., Gundlach, H., Haberer, G., Hollister, J. D., Ossowski, S., Ottilar, R. P., Salamov, A. A., Schneeberger, K., Spannagl, M., Wang, X., Yang, L., … Guo, Y.-L. (2011). The Arabidopsis lyrata genome sequence and the basis of rapid genome size change. Nature Genetics, 43(5), 476–481. 10.1038/ng.807

J, M. H. (1940). Bearing of the Drosophila work on systematics. The New Systematics, 185–268.

Leppälä, J., Bokma, F., & Savolainen, O. (2013). Investigating incipient speciation in Arabidopsis lyrata from patterns of transmission ratio distortion. Genetics, 194(3), 697–708. 10.1534/genetics.113.152561

Luo, X., Xu, N., Huang, J., Gao, F., Zou, H., Boudsocq, M., Coaker, G., & Liu, J. (2017). A Lectin Receptor-Like Kinase Mediates Pattern-Triggered Salicylic Acid Signaling. Plant Physiology, 174(4), 2501–2514. 10.1104/pp.17.00404

Martin, M., Patterson, M., Garg, S., Fischer, S. O., Pisanti, N., Klau, G. W., Schöenhuth, A., & Marschall, T. (2016). WhatsHap: Fast and accurate read-based phasing (p. 085050). bioRxiv. 10.1101/085050

Matute, D. R., & Coyne, J. A. (2010). INTRINSIC REPRODUCTIVE ISOLATION BETWEEN TWO SISTER SPECIES OF DROSOPHILA. Evolution, 64(4), 903–920. 10.1111/j.1558-5646.2009.00879.x

Mirchandani, C. D., Shultz, A. J., Thomas, G. W. C., Smith, S. J., Baylis, M., Arnold, B., Corbett-Detig, R., Enbody, E., & Sackton, T. B. (2023). A fast, reproducible, high-throughput variant calling workflow for evolutionary, ecological, and conservation genomics (p. 2023.06.22.546168). bioRxiv. 10.1101/2023.06.22.546168

Monnet, F., Postel, Z., Touzet, P., Fraïsse, C., Peer, Y. V. de, Vekemans, X., & Roux, C. (2023). Rapid establishment of species barriers in plants compared to animals (p. 2023.10.16.562535). bioRxiv. 10.1101/2023.10.16.562535

Muller, H. J. (1942). Isolating mechanisms, evolution, and temperature. Biol. Symp., 6, 71.

Muralla, R., Lloyd, J., & Meinke, D. (2011). Molecular Foundations of Reproductive Lethality in Arabidopsis thaliana. PLOS ONE, 6(12), e28398. 10.1371/journal.pone.0028398

Noble, L. M., Yuen, J., Stevens, L., Moya, N., Persaud, R., Moscatelli, M., Jackson, J. L., Zhang, G., Chitrakar, R., Baugh, L. R., Braendle, C., Andersen, E. C., Seidel, H. S., & Rockman, M. V. (2021). Selfing is the safest sex for Caenorhabditis tropicalis. eLife, 10, e62587. 10.7554/eLife.62587

Novikova, P. Y., Hohmann, N., Nizhynska, V., Tsuchimatsu, T., Ali, J., Muir, G., Guggisberg, A., Paape, T., Schmid, K., Fedorenko, O. M., Holm, S., Säll, T., Schlötterer, C., Marhold, K., Widmer, A., Sese, J., Shimizu, K. K., Weigel, D., Krämer, U., … Nordborg, M. (2016). Sequencing of the genus Arabidopsis identifies a complex history of nonbifurcating speciation and abundant trans-specific polymorphism. Nature Genetics, 48(9), 1077–1082. 10.1038/ng.3617

Orr, H. A. (1996). Dobzhansky, Bateson, and the Genetics of Speciation. Genetics, 144(4), 1331–1335.

Price, T. a. R., Verspoor, R., & Wedell, N. (2019). Ancient gene drives: An evolutionary paradox. Proceedings of the Royal Society B: Biological Sciences, 286(1917), 20192267. 10.1098/rspb.2019.2267

Ravinet, M., Faria, R., Butlin, R. K., Galindo, J., Bierne, N., Rafajlović, M., Noor, M. A. F., Mehlig, B., & Westram, A. M. (2017). Interpreting the genomic landscape of speciation: A road map for finding barriers to gene flow. Journal of Evolutionary Biology, 30(8), 1450–1477. 10.1111/jeb.13047

Roux, C., Castric, V., Pauwels, M., Wright, S. I., Saumitou-Laprade, P., & Vekemans, X. (2011). Does speciation between Arabidopsis halleri and Arabidopsis lyrata coincide with major changes in a molecular target of adaptation? PloS One, 6(11), e26872. 10.1371/journal.pone.0026872

Roux, C., Fraïsse, C., Romiguier, J., Anciaux, Y., Galtier, N., & Bierne, N. (2016). Shedding Light on the Grey Zone of Speciation along a Continuum of Genomic Divergence. PLOS Biology, 14(12), e2000234. 10.1371/journal.pbio.2000234

Seymour, D. K., Chae, E., Arioz, B. I., Koenig, D., & Weigel, D. (2019). Transmission ratio distortion is frequent in Arabidopsis thaliana controlled crosses. Heredity, 122(3), 294–304. 10.1038/s41437-018-0107-9

Simon, A., Bierne, N., & Welch, J. J. (2018). Coadapted genomes and selection on hybrids: Fisher’s geometric model explains a variety of empirical patterns. Evolution Letters, 2(5), 472–498. 10.1002/evl3.66

Stein, R. J., Höreth, S., de Melo, J. R. F., Syllwasschy, L., Lee, G., Garbin, M. L., Clemens, S., & Krämer, U. (2017). Relationships between soil and leaf mineral composition are element-specific, environment-dependent and geographically structured in the emerging model Arabidopsis halleri. New Phytologist, 213(3), 1274–1286. 10.1111/nph.14219

Turelli, M., & Orr, H. A. (1995). The dominance theory of Haldane’s rule. Genetics, 140(1), 389–402. 10.1093/genetics/140.1.389

Wang, W.-K., Ho, C.-W., Hung, K.-H., Wang, K.-H., Huang, C.-C., Araki, H., Hwang, C.-C., Hsu, T.-W., Osada, N., & Chiang, T.-Y. (2010). Multilocus analysis of genetic divergence between outcrossing Arabidopsis species: Evidence of genome-wide admixture. The New Phytologist, 188(2), 488–500. 10.1111/j.1469-8137.2010.03383.x

Wang, Y., Cordewener, J. H. G., America, A. H. P., Shan, W., Bouwmeester, K., & Govers, F. (2015). Arabidopsis Lectin Receptor Kinases LecRK-IX.1 and LecRK-IX.2 Are Functional Analogs in Regulating Phytophthora Resistance and Plant Cell Death. Molecular Plant-Microbe Interactions®, 28(9), 1032–1048. 10.1094/MPMI-02-15-0025-R

Weichenhan, D., Traut, W., Kunze, B., & Winking, H. (1996). Distortion of Mendelian recovery ratio for a mouse HSR is caused by maternal and zygotic effects. Genetical Research, 68(2), 125–129. 10.1017/s0016672300034017

Xiong, Y., Contento, A. L., & Bassham, D. C. (2005). AtATG18a is required for the formation of autophagosomes during nutrient stress and senescence in Arabidopsis thaliana. The Plant Journal, 42(4), 535–546. 10.1111/j.1365-313X.2005.02397.x

