## Supplementary figures and images for "Diverging Arabidopsis populations quickly accumulate pollen-acting genetic incompatibilities"

### Supplemental Figure 2

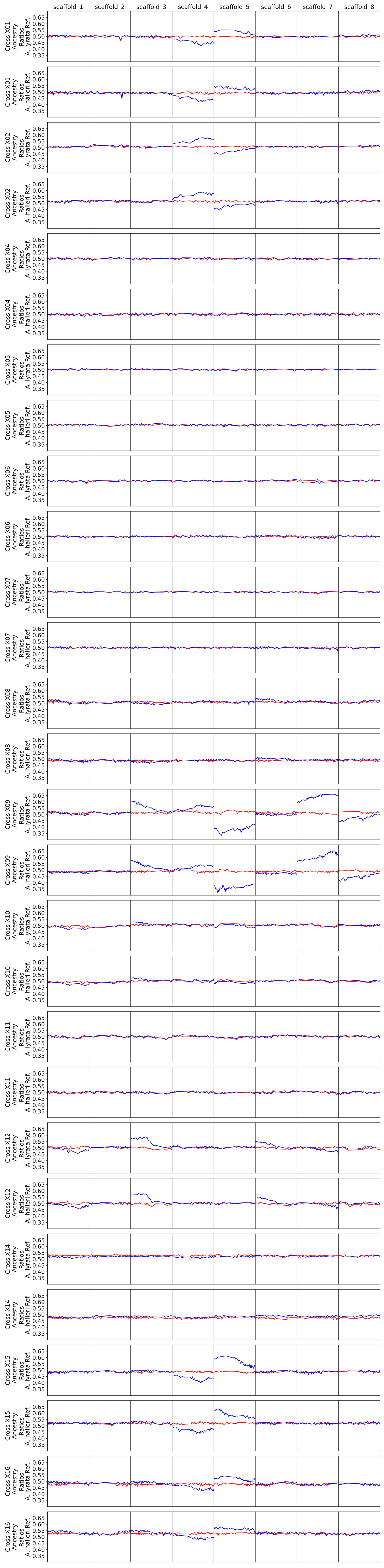
